# State of mangrove biodiversity assessment in Kenya and the prospect of environmental DNA in strengthening surveys

**DOI:** 10.64898/2026.03.14.711771

**Authors:** Oketch Fredrick Onyango, Judith Okello, Zipporah Muchiri, Samuel Mwamburi, Chepkemboi Labatt, Everline Osir Owiro, Shawlet Cherono

**Affiliations:** Kenya Marine and Fisheries Research Institute, P.O. Box 81651 – 80100, Kenya; Department of Biochemistry and Biotechnology, School of Pure and Applied Sciences, Pwani University, P. O. Box 195-80108, Kilifi, Kenya; Department of Biochemistry and Molecular Biology, Faculty of Science, Egerton University, P.O. Box 536-20115, Egerton-Njoro, Kenya; Department of Medicinal Plants Research and Drug Development, PAULESI, P.O. Box 22133, University of Ibadan, Ibadan, Oyo State, Nigeria; Department of Earth and Environmental Sciences, KU Leuven, Belgium

**Keywords:** Environmental DNA, Mangrove ecosystem, Biodiversity assessment, Taxonomic coverage, Kenya

## Abstract

Assessing and monitoring biodiversity in mangrove ecosystems remains challenging, with most studies relying on proxy indicators to infer biodiversity status. This limit understanding of biodiversity dynamics and constrains evidence-based mangrove management. In the Western Indian Ocean region, biodiversity assessments in mangrove forests remain scanty, with no clear information on spatiotemporal and taxonomic coverage. Addressing these gaps requires examining existing biodiversity records and exploring complementary approaches that can broaden the scope and efficiency of biodiversity monitoring. This study assessed the current state of biodiversity assessments in mangrove forests in Kenya and evaluated the feasibility of environmental DNA (eDNA) as a complementary biodiversity monitoring tool. A systematic literature review was conducted by retrieving published sources from major academic databases using defined search terms to extract and compile taxonomic information. In addition, a snapshot eDNA survey was carried out in selected mangrove forests, where sediment and water samples were collected, processed, and analyzed using established molecular and bioinformatics pipelines. The literature review identified 26 sources documenting biodiversity across 15 mangrove forest areas, with 68% of the studies concentrated in four sites representing about 6% of mangrove cover in Kenya. A total of 1,044 unique taxa belonging to 255 families were identified, with the classes *Teleostei, Aves, Chromadorea,* and *Malacostraca* accounting for 84.5% of documented taxa. The eDNA survey detected heterogeneous taxa from multiple ecosystems, including 502 taxa belonging to 305 families. Only 67 families were common to both datasets, highlighting the complementarity of literature-based inventories and eDNA detection. While eDNA showed considerable potential to expand biodiversity detection, its application is constrained by a number of factors. Integrating eDNA as a core biodiversity monitoring tool in mangroves will require combining conventional surveys with molecular tools, developing curated regional DNA reference databases, and adopting standardized analytical frameworks.

## Introduction

Mangrove ecosystems are among the most productive and ecologically important coastal habitats, providing ecosystem goods and services that sustain biodiversity and support the livelihoods of millions of people in tropical coastal regions (Ahmed et al., 2022; Ayyam et al., 2019; Huxham et al., 2017; Olima et al., 2025). Although mangrove forests are characterized by relatively low diversity of higher plant species (Spalding et al., 2010; FAO, 2023), they support remarkable diversity of other life forms and function as breeding, foraging, and nursery habitats for fish and invertebrates (Arceo-Carranza et al., 2021; Das et al., 2022). Biodiversity patterns within mangrove ecosystems are shaped by complex interactions among abiotic and biotic factors (Lovelock et al., 2025). However, anthropogenic pressures and climate-related stressors increasingly influence the structure and composition of mangrove-associated communities (Alongi, 2022; Bosire et al., 2014; Liu et al., 2024; Tasnim et al., 2023; Carugati et al., 2018), with consequential effects on ecosystem functioning and service delivery (Hülsen et al., 2025). In response to these pressures, conservation initiatives, primarily through ecological restoration and the implementation of nature-based solutions, are being adopted to reduce stress on mangrove resources (Kinya et al., 2024; Lovelock et al., 2024; Mahmood et al., 2023). However, as observed in many ecosystems in tropical developing regions, systematic and comprehensive biodiversity assessments remain limited (Collen et al., 2008; Friess, 2025; Guerrero-Casado & Monge-Nájera, 2021; Moreno et al., 2023; Titley et al., 2017; Tydecks et al., 2018). This lack of coordinated biodiversity monitoring makes it challenging to effectively integrate biodiversity data into mangrove management and conservation frameworks.

In Kenya, mangrove forests account for approximately 3% of the national forest cover (GoK, 2017). A substantial proportion of these forests, estimated at 40.1%, is considered ecologically degraded (Bosire et al., 2014; GoK, 2017; Hamza et al., 2022), while biodiversity data across much of their distribution remain limited. Persistent gaps in biodiversity monitoring constrain large-scale ecological inference and may lead to biased assessments (Bowler et al., 2025). In the absence of standardized biodiversity monitoring frameworks, uncertainties remain regarding the indicators currently used to evaluate recovery processes after management or restoration interventions (Andradi-Brown et al., 2013; Dalton et al., 2023). Addressing these limitations requires the redesign of monitoring frameworks through the adoption of techniques capable of generating comprehensive biodiversity data (Orr et al., 2022). Current knowledge of mangrove biodiversity in Kenya derives largely from site-specific, sporadic, and taxon-focused surveys that, while informative, present fragmented information and fail to capture the spatiotemporal and ecological variability characteristic of these ecosystems (Gonzalez et al., 2023). The adoption of scalable and standardized assessment approaches would improve data comparability across sites and over time, thus enabling a more integrated understanding of biodiversity patterns and ecosystem processes (Leit et al., 2025; Makiola et al., 2020; McGlone et al., 2020). Without such methodological advances, biodiversity knowledge gaps are likely to persist, constraining effective ecosystem management and limiting the capacity to track progress toward global conservation targets such as Sustainable Development Goals 14 and 15, the Convention on Biological Diversity, and the post-2020 Global Biodiversity Framework.

Conventional biodiversity survey methods have played an important role in establishing baseline information and documenting specific taxonomic groups in selected mangrove forests (Gajdzik *et al*., 2014; Kimani *et al*., 1997; Nyunja *et al*., 2009; Waweru *et al*., 2022). However, their broader application has been constrained by several factors, limited capacity and expertise (Stephenson *et al*., 2020). Environmental DNA (eDNA) has emerged as a scalable and cost-effective alternative to conventional survey techniques, enabling non-invasive detection of a broad range of taxonomic groups (Schwentner *et al*., 2021). Although the persistence of eDNA in environmental matrices varies with local conditions (Joseph *et al*., 2022; Nielsen *et al*., 2007; Stewart, 2019), this variability allows eDNA to capture contemporary community composition and, in relatively undisturbed systems, detect recent past biodiversity (Ariza *et al*., 2023; Sullivan *et al*., 2025). These attributes make eDNA particularly well-suited for biodiversity assessments in complex and sensitive ecosystems such as mangroves.

Reflecting on its growing utility, the application of eDNA in ecological research has expanded rapidly over the past decade, increasing from only a handful of studies in 2010 to several hundred conducted annually by the early to mid-2020s, indicating widespread adoption across diverse ecosystems (Sahu *et al*., 2025). Despite this growth, the use of eDNA in mangrove forests in Kenya has so far been largely confined to the assessment of microbial assemblages (Muwawa *et al*., 2021, 2024), leaving its potential for comprehensive, multi-taxa biodiversity assessment largely unexplored. Against this background, key questions remain regarding the current state of documented taxa, the extent of spatiotemporal and taxonomic coverage in existing studies, and the feasibility of eDNA to streamline biodiversity assessment in mangroves in Kenya. To address these questions, this study evaluated the progress of biodiversity assessment in Kenya over the years and assessed the utility of eDNA as a complementary tool for biodiversity assessment, providing a synthesis of mangrove biodiversity research in Kenya, identifying existing knowledge gaps, and evaluating the feasibility of eDNA in strengthening biodiversity surveys.

## Materials and methods

### Study area

The Kenyan coastline covers more than 500 km, extending from Ishakani (1°39’ S, 41°33’ E) near the Kenya-Somalia border to Jimbo (4°40’ S, 39°13’ E), near the Kenya-Tanzania border (Kirui *et al*., 2013). Characterised by a range of physical features and varying anthropogenic activities which have shaped the dynamics of biodiversity within its bounds. At the marine-terrestrial interface are mangrove forests distributed in a disjunct pattern across estuaries, bays, and archipelagos. The forests cover an estimated 61,271 hectares, of which approximately 24,585 hectares have been lost (GoK, 2017). The largest proportion of mangrove forest cover in Kenya is found in Lamu County (61%), followed by Kwale and Kilifi Counties (14% each), with smaller proportions located in Mombasa (6%) and Tana River Counties (5%) (GoK, 2017). In this study, a systematic literature review targeted all known mangrove forest formations in Kenya. Meanwhile, the environmental DNA (eDNA) survey was conducted in selected mangrove forests of Vanga, Mwena, Kongo River, Sii Island, Mtwapa Creek, Takaungu Creek, Kilifi Creek, Marereni, and Lamu Southern Swamp, all of which are located across different regions along the Kenyan coast (Fig. 1).

**Figure 1.**
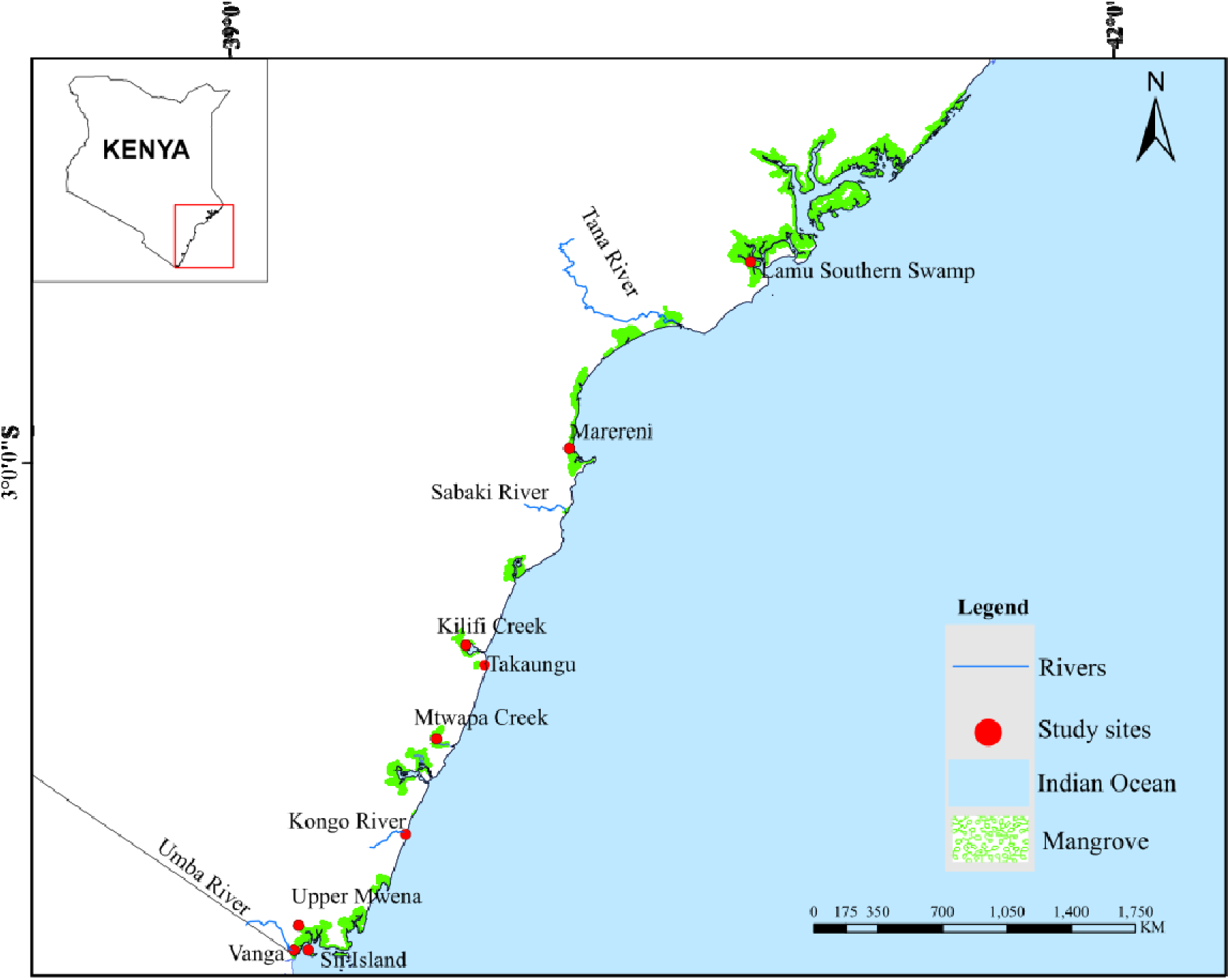
Sampling locations for the eDNA survey. Red markers indicate eDNA sample collection sites, while green polygons represent mangrove forest blocks along the Kenyan coast.

### Literature review, data curation and analysis

Literature search was conducted between October and December 2024, following the guidelines outlined in the Preferred Reporting Items for Systematic Reviews and Meta-Analyses (PRISMA) (Moher *et al*., 2009). A hybrid search criterion was used, combining a structured systematic search of peer-reviewed literature in the Web of Science, Scopus, and ProQuest databases, as illustrated by (Testa *et al*., 2025), combined with a manual (hands-on) search for grey literature in the Google Scholar database following the approach illustrated by (Haddaway *et al*., 2015). All searches were limited to biodiversity surveys within the mangrove forest in Kenya, with no restrictions placed on the date of publication. A standardized search string ("Biodiversity" OR "Biodiversity status" OR "Biodiversity survey" OR "Biodiversity assessment" OR "Taxon survey") AND ("Mangrove*") AND ("Kenya") NOT ("Coral*" OR "Seagrass*" OR "Terrestrial coastal forest*") was used across all the aforementioned databases. Literature sources with relevant information were retrieved, compiled in a spreadsheet database, and manually screened for information. Of the 156 sources identified, 26 had information on biodiversity surveys in mangroves in Kenya. Taxonomic data and relevant metadata were extracted and compiled in a spreadsheet. The taxa list generated (herein referred to as Checklist) was matched in the World Register of Marine Species (WoRMS) using worrms package v0.4.3 (Chamberlain & Vanhoorne, 2023) in R. Only taxa matched correctly to genus levels were retained and parsed for downstream analysis in R v4.5.1 (R Core Team, 2024) using a suite of tidyverse packages (Wickham *et al*., 2019). Potential bias in biodiversity studies across mangrove forests and taxonomic groups was evaluated using Gini coefficient.

### Environmental DNA (eDNA) sample collection

Samples were collected following standard eDNA sampling protocols outlined by Minamoto *et al*. (2021). Water samples were collected during high tide along tidal channels, following the sampling design described by Bruce *et al*. (2021). At each site, at approximately 50-meter intervals, 25 sub-samples of 200 ml were collected from the water surface at multiple points and then pooled into a sterile 5,000 ml container (Olympus Lyse Bottle, China) to make a composite water sample. Triplicate composite water samples of 5,000 ml were collected at each site, ensuring representative sampling in all mangrove inundation classes. For sediment samples, three representative 30 m × 30 m plots were established per site. At each plot, surface sediments (∼2 cm deep) were randomly sampled using single-use sterile spatulas and then pooled into sterile 50 ml Falcon tubes. Triplicate samples were collected from each plot. GPS coordinates were recorded for each sampling location. To minimize cross-contamination, water and sediment samples were stored in separate compartments of a cooler box, and each site was sampled on a separate day using new sterile equipment for every field round to prevent cross-site contamination. All samples were transported to the laboratory within one hour of collection and temporarily stored at -80°C for no more than one day prior to water filtration and eDNA extraction.

### Sample processing and DNA extraction

Prior to sample processing and eDNA extraction, all reusable materials and working surfaces were sterilised following Goldberg *et al*., (2016). Samples were retrieved from the freezer and allowed to thaw at room temperature (27°C). The external surfaces of the containers were then wiped with 50% bleach, followed by 70% ethanol. The water filtration unit (Vacuum Manifold kit, Perkin Elmer, USA) was assembled in a laminar flow hood (Aelous V, Japan), and UV-sterilised for 30 minutes. Using 0.22 μm nitrocellulose filters (Scharlab, Spain), water was filtered for each study site, generating a varied number of filter replicates. DNA was extracted from the sediment and nitrocellulose filters using ZymoBIOMICS™ DNA Miniprep Kit (Zymo Research, USA), incorporating the bead homogenization step as described in the manufacturer’s protocol. All the extracted DNA were eluted to a final volume of 50 μL using ZymoBIOMICS™ DNase/RNase-free water. The purity and concentration of the DNA were assessed using a NanoDrop spectrophotometer (Thermo Scientific, USA). All the 78 samples were processed and sent to a sequencing service provider (Inqaba Biotec, South Africa) for library preparation and high-throughput sequencing following the Illumina MiSeq library preparation and sequencing protocol.

### Bioinformatics analysis and taxonomic assignments

Raw shotgun sequences were first assessed for quality using FastQC (Brown *et al*., 2017), and the results were aggregated using MultiQC v1.23 (Ewels *et al*., 2016) to provide an overview of sequence quality across all samples. Low quality reads and adapters were removed using Fastp v0.23.4 (Chen *et al*., 2018) with parameters set as: reads with quality score below 20 (-q 20), and/or have more than 30% of bases falling below the quality threshold (-u 30) were removed; sequences containing more than 5 ambiguous bases (-n 5) were filtered and removed; and only reads with a minimum length of 50 base pairs (--length_required 50) were retained. Reads that passed these filters were *de novo* assembled into contigs using SPAdes v3.15.5 (Prjibelski *et al*., 2020), applying the metaSPAdes option to optimize assembly for metagenomic sequences. An additional filtering step was implemented where contigs shorter than 200 base pairs were discarded. For taxonomic classification, the assembled contigs were queried against a Kraken2 database customised from the National Center for Biotechnology Information RefSeq database (https://www.ncbi.nlm.nih.gov/refseq/, accessed on January 30, 2025). The database included taxonomically indexed genomes alongside corresponding NCBI taxonomy files. Kraken2 was executed with the *--use-names* flag to produce taxonomic outputs at 97% sequence similarity. The resulting classification reports were processed in Pavian (Breitwieser & Salzberg, 2020) to produce an appropriate file format for downstream analysis.

### Data processing and validation against the regional biodiversity checklist

The Operational Taxonomic Unit (OTU) list generated from the bioinformatics analysis was first matched against the Global Biodiversity Information Facility (GBIF; https://www.gbif.org/) taxonomic backbone using the rgbif package v3.8.1 (Chamberlain *et al*., 2017). Only OTUs with an exact match at the genus level were retained for downstream analyses. The retained OTUs were then cross-referenced with a regional (Western Indian Ocean) biodiversity checklist compiled from online databases and literature, following the guidelines outlined by Jacquemot *et al*. (2024). OTUs present in the eDNA dataset but absent from the regional checklist were labeled “Unexpected”, whereas those present in both the eDNA dataset and the checklist were labeled “Expected” (Jacquemot *et al*., 2024). To obtain habitat information, OTUs were queried against the World Register of Marine Species (https://www.marinespecies.org/) using the function *wm_records_names* in worrms package v0.4.3 (Chamberlain & Vanhoorne, 2023) in R to retrieve complete records including habitat information for each of the detected OTUs. Where automated retrieval was unsuccessful, habitat information was manually extracted from the IUCN Red List database (https://www.iucnredlist.org/), FishBase (https://www.fishbase.se/), and Ocean Biodiversity Information System (OBIS; https://obis.org/). OTUs of Bacteria, Fungi, and Archaea known to inhabit multiple environments were classified as ubiquitous. All non-marine OTUs (except those from brackish environments and those classified as ubiquitous), unexpected, as well as OTUs with low relative abundance (< 0.0001) and low occurrence frequency (< 0.1), were excluded to reduce sequencing artefacts, minimize data sparsity, and improve interpretability of downstream diversity and community analyses. Transformation and initial analysis were performed using the microeco package v1.16.0 (Liu *et al*., 2021) in R v4.4.2 (R Core Team, 2024).

### Diversity indices calculation and statistical analysis

All statistical analyses were performed in R v4.4.2 (R Core Team, 2024). To meet chi-square test assumptions, sites with low taxa counts were pooled into a single category prior to analysis. Alpha diversity indices (Chao1 richness, Shannon diversity, and Simpson diversity) were calculated using the vegan package v2.7-1 (Oksanen *et al*., 2013). Differences in community composition between sampling sites and sample types were assessed using Permutational Multivariate Analysis of Variance (PERMANOVA) with 999 permutations, implemented using the adonis2 function in vegan. In addition, Analysis of Similarities (ANOSIM) was performed in vegan with 999 permutations to provide an independent, rank-based test of community separation that is less sensitive to variation in within-group dispersion. Community patterns were visualized using non-metric multidimensional scaling (nMDS) based on Bray-Curtis dissimilarities, calculated using the metaMDS function in vegan (k = 2, trymax = 100). To identify OTU families contributing most to differences in beta diversity, similarity percentage analysis (SIMPER) was performed using the simper function in vegan.

## Results

### Overview of the literature review data

Using the defined search criteria, a total of 26 studies on biodiversity surveys across mangrove forests in Kenya were identified. These studies were published between 1990 and 2024 (Fig. S1, Table S1), with the majority (66.67%) being peer-reviewed journal articles (Fig. 4). A large proportion of the studies (92.59%) were based on primary data collection using a range of conventional survey methods and, more recently, environmental DNA-based techniques (Fig. 4; Fig. S2). Three additional studies, published in 1996, 2010, and 2021, relied on secondary data sources to synthesize the general biodiversity in coastal forest in Kenya, avian diversity in Mida Creek, and mangrove biodiversity in Gazi Bay, respectively (Table S1). The number and frequency of the studies varied across the surveyed sites. Significant variation was identified in publication output (χ² test, df =25, p = 0.004), with survey effort disproportionately skewed towards selected mangrove forests (Gini coefficient = 0.50). Most of the studies (68.56%) focused on four mangrove forests, viz. Gazi Bay, Mida Creek, Tudor Creek, and Kilifi Creek. Gazi Bay was the most frequently studied site, accounting for more than a third (34.2%) of the biodiversity survey frequency, followed by Mida Creek (21.1%), Tudor Creek (7.9%), and Kilifi Creek (5.3%). Other sites, that is, Bamburi, Kanamai, Mkomani, and Malindi, all surveyed in 1990, Sabaki (1995), Shimoni (1998), Mtwapa Creek (2004), Ungwana Bay (2010), and Vanga (2022), survey were only conducted once and were not featured in any subsequent studies (Fig. S3a). Over the years, publication outputs increased marginally (p = 0.897, R² < 0.01, Fig. 2a), averaging three studies in four years, with peaks of three studies in 2009 and 2010 (Fig. S1). None of the reviewed studies was linked to ongoing biodiversity monitoring programmes or to any online biodiversity repositories.

**Figure 2.**
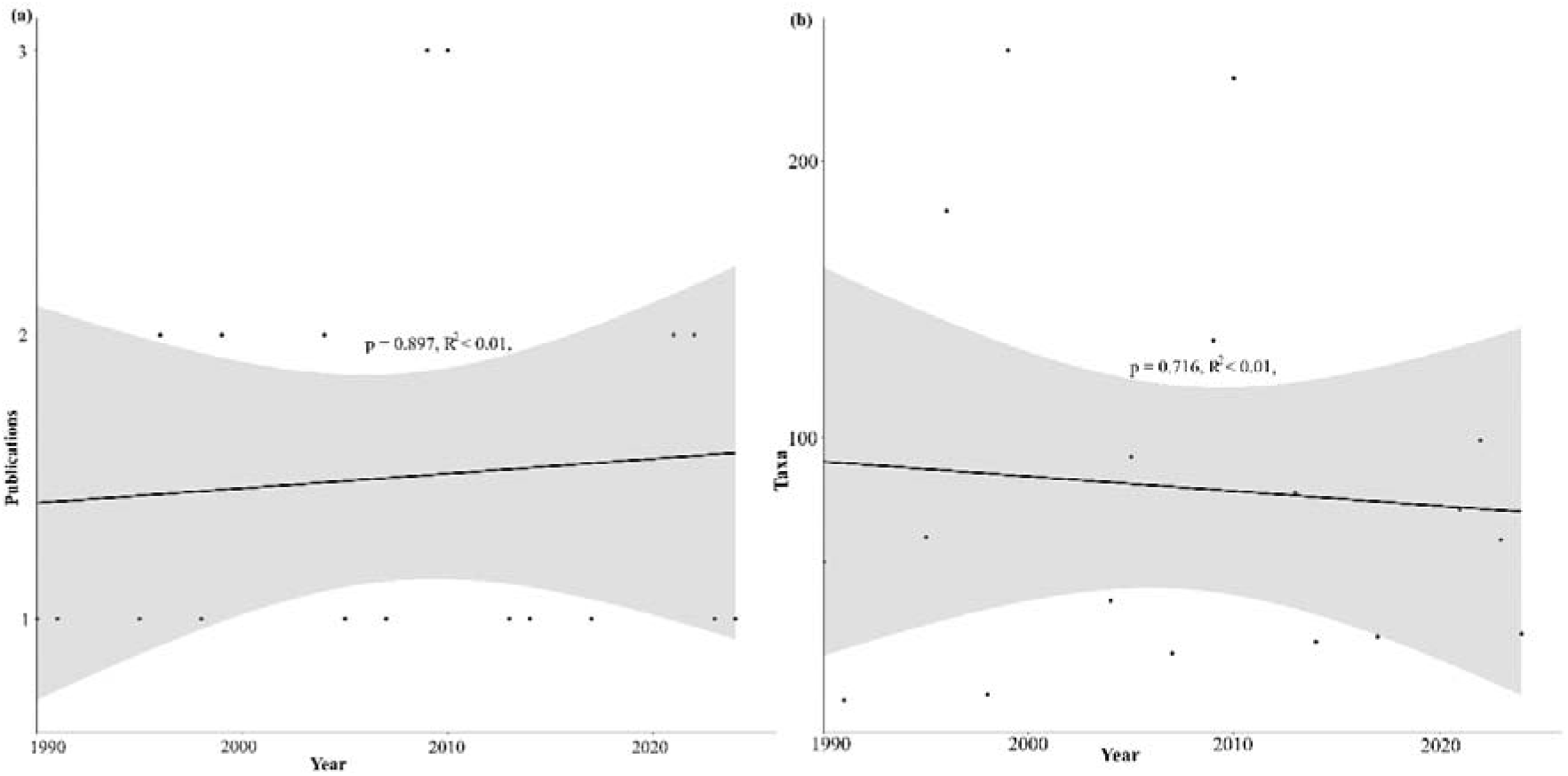
Trends in mangrove biodiversity assessments in Kenya. (a) Temporal trend in publications on mangrove biodiversity since 1990. (b) Temporal trend in the number of unique taxa surveyed in mangrove forests in Kenya since 1990.

### Taxa retrieved from the reviewed documents

The diversity of taxa recorded, evaluated at class level, over the years varied across the mangrove forests (Kruskal-Wallis χ² = 11,189, df = 28, p < 2.2 × 10L¹L), with cumulative taxa survey fluctuating and showing no evidence of a sustained increase in coverage over the years (Mann-Kendall τ = 0.046, p = 0.820). As a result, many studies repeatedly documented previously recorded taxa, contributing marginally to the recording of new taxonomic groups (Mann-Kendall τ = -0.052, p = 0.805, Fig. 3b). In total, 1,044 distinct taxa spanning 29 classes, and 255 families were retrieved from the studies. Gazi Bay had the highest and most diverse documented taxa belonging to 6 kingdoms, 19 classes, and 143 families, followed by Mida Creek, where taxa from 5 kingdoms, 16 classes, and 135 families were recorded. Sites such as Sabaki, Mtwapa Creek, Shimoni, Kanamai, Mkomani, and Bamburi had lower records of biodiversity, with than six classes of taxa identified (Table S2). In studies categorised as general (nonspecific to any mangrove forest block and mostly compiled from secondary sources) a range of taxa belonging to 2 kingdoms, 9 classes, and 83 families were retrieved (Table S2). Temporally, there were no notable increase in the number of taxa surveyed (Fig. 2b; p = 0.716, R² < 0.01). Among the identified taxa, there was a disproportionate focus on *Teleostei, Aves, Malacostraca*, and *Chromadorea* (Fig. 3a; Fig. S6). The class *Teleostei* was the most represented and was consistently recorded over the years (Fig. 3c), accounting for 56.13% of all unique taxa retrieved from literature sources (Fig. 4). Other classes with significant representation in the literature review were *Aves* (17.34%), *Chromadorea* (7.18%), *Malacostraca* (4.79%), *Mammalia* (2.58%), *Enoplea* (2.59%), *Magnoliopsida* (2.30%), and *Gastropoda* (1.63%). The remaining classes accounted for 1% (≤ 8 taxa) of all the taxa in the literature review (Fig. 4).

**Figure 3.**
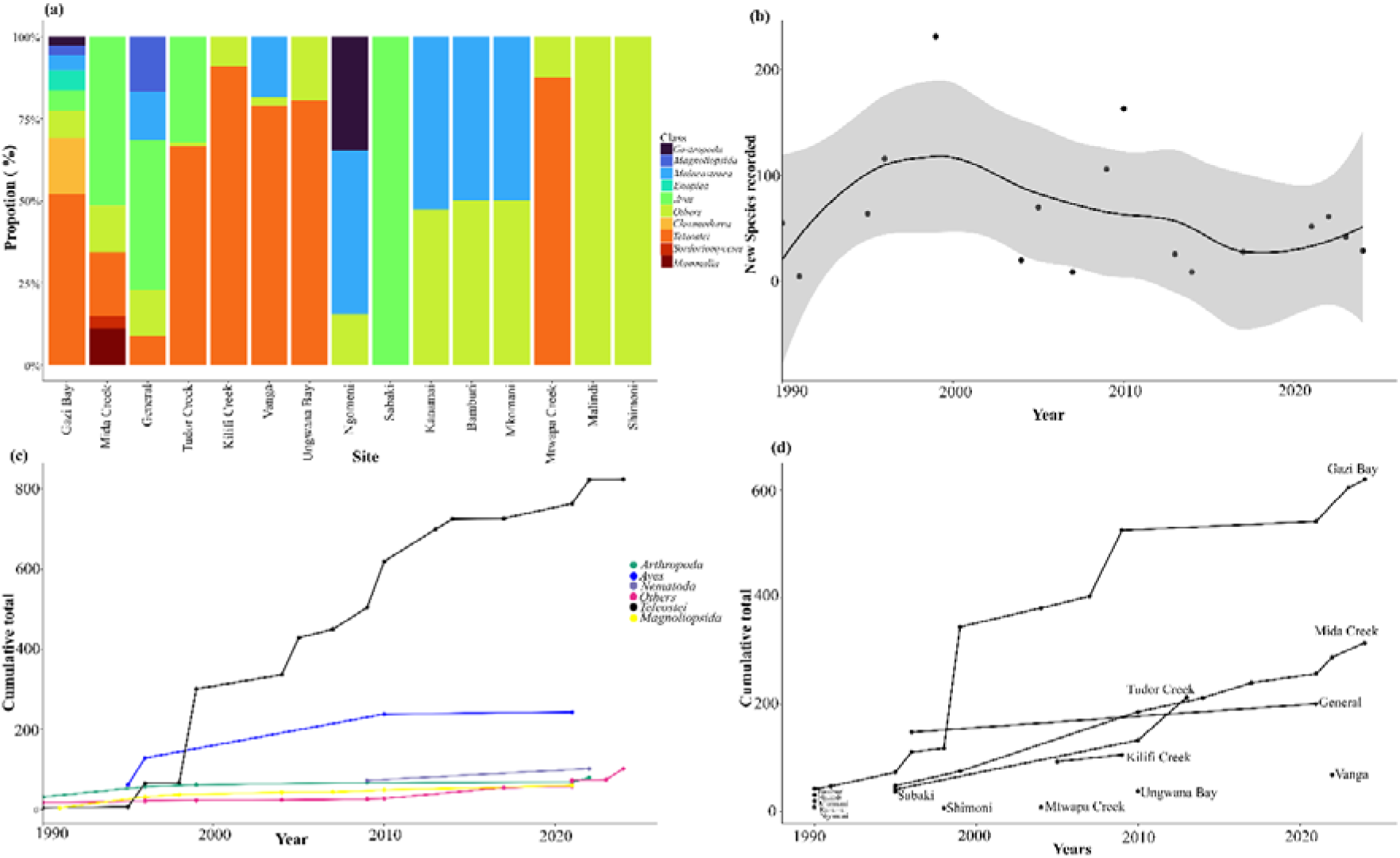
Taxonomic profile of biodiversity records identified in the literature review. (a) Distribution of taxa (class level) recorded across sampling sites. (b) Temporal trend in the number of species surveyed. (c) Cumulative total number of taxa documented over the years. (d) Cumulative total of taxa surveyed at each site over time

**Figure 4.**
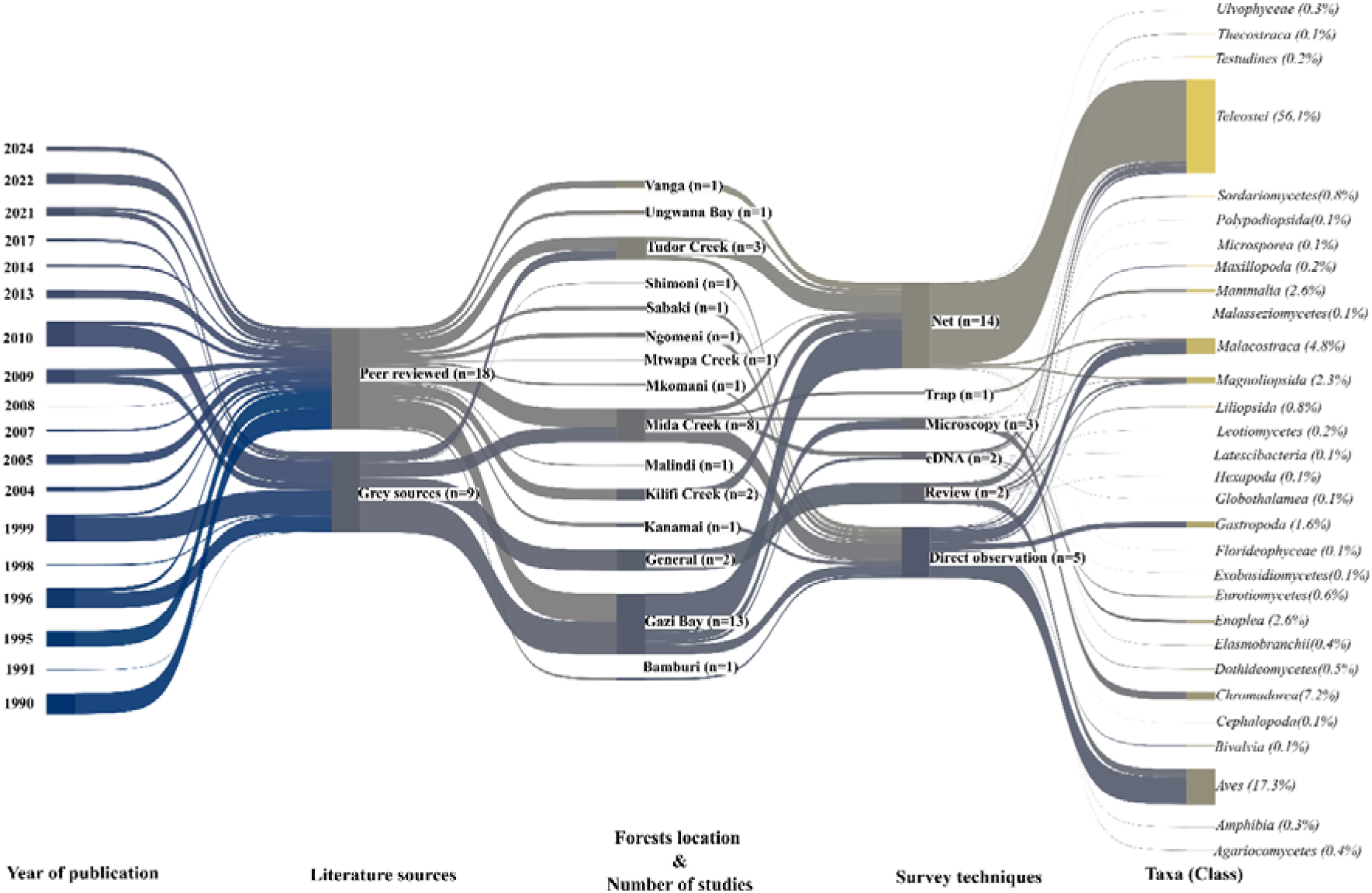
Sankey diagram summarizing key outputs from reviewed studies on the status of biodiversity assessments in mangrove forests in Kenya over time. The diagram illustrates linkages among study year, document type in which survey outputs were reported, mangrove forests surveyed, survey techniques used, and the taxa identified (classified to class level). Some sites were reported in multiple studies; therefore, the frequency of sites (n = 38) exceeds the number of reviewed publications (n = 26). Node size is proportional to the number of species recorded.

### Techniques used in the historical surveys

The taxa list retrived from literature review was largely dependent on the survey techniques (χ² = 17,844.8, df = 364, p < 0.001, Cramér’s V = 0.508). Field-based methods favored the survey of conspicuous and easy-to-observe taxa, while the laboratory techniques were used to identify microscopic organisms (Fig. S2). Use of various types of nets was the most common approach, mainly in the survey of *Teleostei, Cephalopoda*, and *Elasmobranchii* (Fig. S3b; Fig. S2). Observation was used in the assessment of *Malacostraca, Aves, Bivalvia, Florideophyceae, Gastropoda, Thecostraca and Ulvophyceae* (Fig. S2). Microscopy was used predominantly to assess *Nematoda* and *Microsporidia*. For plants, particularly the class *Magnoliopsida*, surveys combined a transect with direct observation. The application of environmental DNA metabarcoding was mainly used in the assessment of bacterial and fungal diversity associated with selected mangrove species (Fig. S2). A review of secondary sources was used in three reports, where a taxa list was compiled from previously published sources. The choice of technique used depended on the taxa targeted in the survey.

### eDNA sequencing results and OTU verification against regional species checklists

Sequencing of 78 sample libraries generated a total of 513,311,730 raw reads (mean = 5,968,741; SD = ±606,140) that were assembled into 5,934,025 contigs (mean = 76,077; SD = ±2,956). From these, 97,843 contigs (mean = 1,254; SD = ±342), representing 1.65% of the assembled contigs, were clustered and assigned to 6,572 OTUs at >=98% sequence similarity. After the removal of OTUs with low absolute sequence abundance and low occurrence frequency, 2,330 OTUs were retained. Of the retained OTUs, 43.81% (n = 995) were putatively identified as originating from terrestrial environments, 29.06% (n = 697) from marine habitats, 14.18% (n = 322) from freshwater systems, and 9.42% (n = 236) were inhabitants of brackish environments. The remaining 3.52% (n = 80) comprised ubiquitous bacterial and fungal lineages (Fig. 5a). Comparison of OTUs against the regional species checklist identified that 36.46% (n = 857) of OTUs were a correct match to species previously recorded in the region (“expected”), whereas 63.54% (n = 1,473) did not match to taxa listed in the checklist (“unexpected”), see Figure 5b. Among the expected OTUs, the ubiquitous *Eubacteriales* was the most represented, followed by Gobiidae, Fabaceae, Muraenidae, and Lutjanidae (Table S3). While the “unexpected” OTUs were dominated by Cyprinidae, Peronosporaceae, Fabaceae, Cicadellidae, Gobiidae, Gracilariaceae, and Bangiaceae (Table S3).

**Figure 5.**
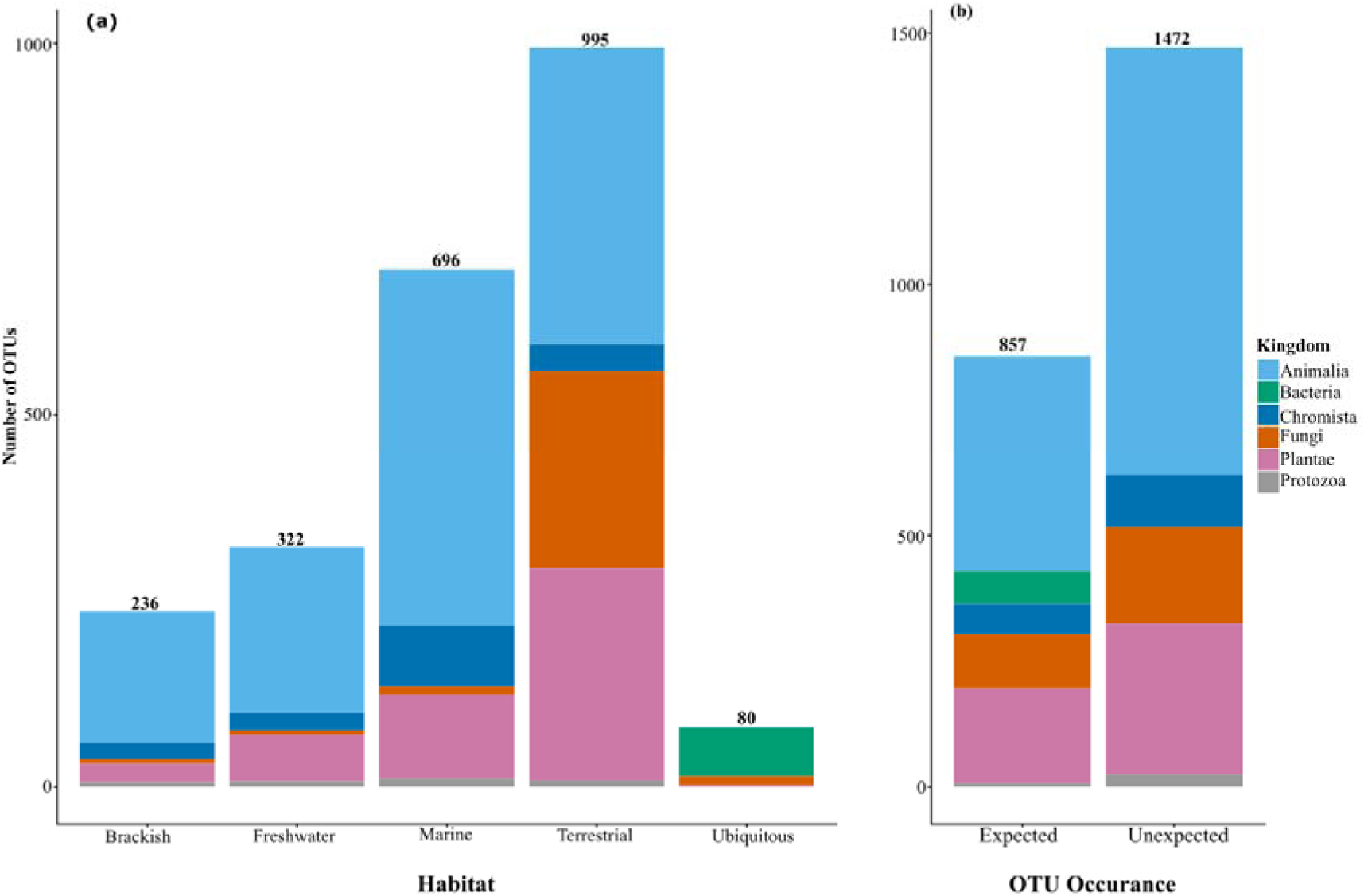
Stacked bar charts illustrating the distribution of operational taxonomic units (OTUs) detected using eDNA. (a) OTUs categorized by inferred habitat of origin. (b) OTUs classified based on regional occurrence records, where “expected” taxa represent species previously documented in the Western Indian Ocean region and “unexpected” taxa represent species not previously recorded in the region.

### Environmental DNA detected marine OTUs and composition

After excluding 1,828 (78.45%) non-marine and “unexpected” OTUs, 502 OTUs belonging to 76 classes, and 305 families were retained (Table S4). Of these, 422 were associated with marine and brackish environments, while 80 comprised ubiquitous bacteria and fungi. Among the OTUs, the kingdom bacteria had the highest contig relative abundance, with the classes *Alphaproteobacteria, Actinomycetota, Betaproteobacteria, Bacteroidota*, and *Gammaproteobacteria* showing high relative abundance across samples (Fig. 6). Other classes, such as *Teleostei, Bacilliota, Ulvophyceae*, and *Deinocota*, also showed relatively high contig abundance (Fig. 6). OTUs from the kingdom *Plantae, Chromista, Fungi,* and *Protozoa* had low contig relative abundance across samples, with most showing cross-sample relative abundance of below 1% (Fig. 6). Based on OTU counts, kingdom Animalia was the most represented, comprising 30 classes with a total of 320 OTUs (Table S4), the majority (63.13%) of which belonged to the class *Teleostei* (Table S5). The kingdom Bacteria was represented by seven classes, with *Bacilliota* accounting for the highest proportion of OTUs, while *Bacteriodonta* and *Deinococcota* accounting for the least (Table S5, Table S7). Plantae comprised 14 classes, dominated by *Florideophyceae*, while *Chromista, Fungi,* and *Protozoa* were represented by 11, 8, and 5 classes, respectively, with varying numbers of OTUs, while protozoa was the least represented (Table S5).

**Figure 6.**
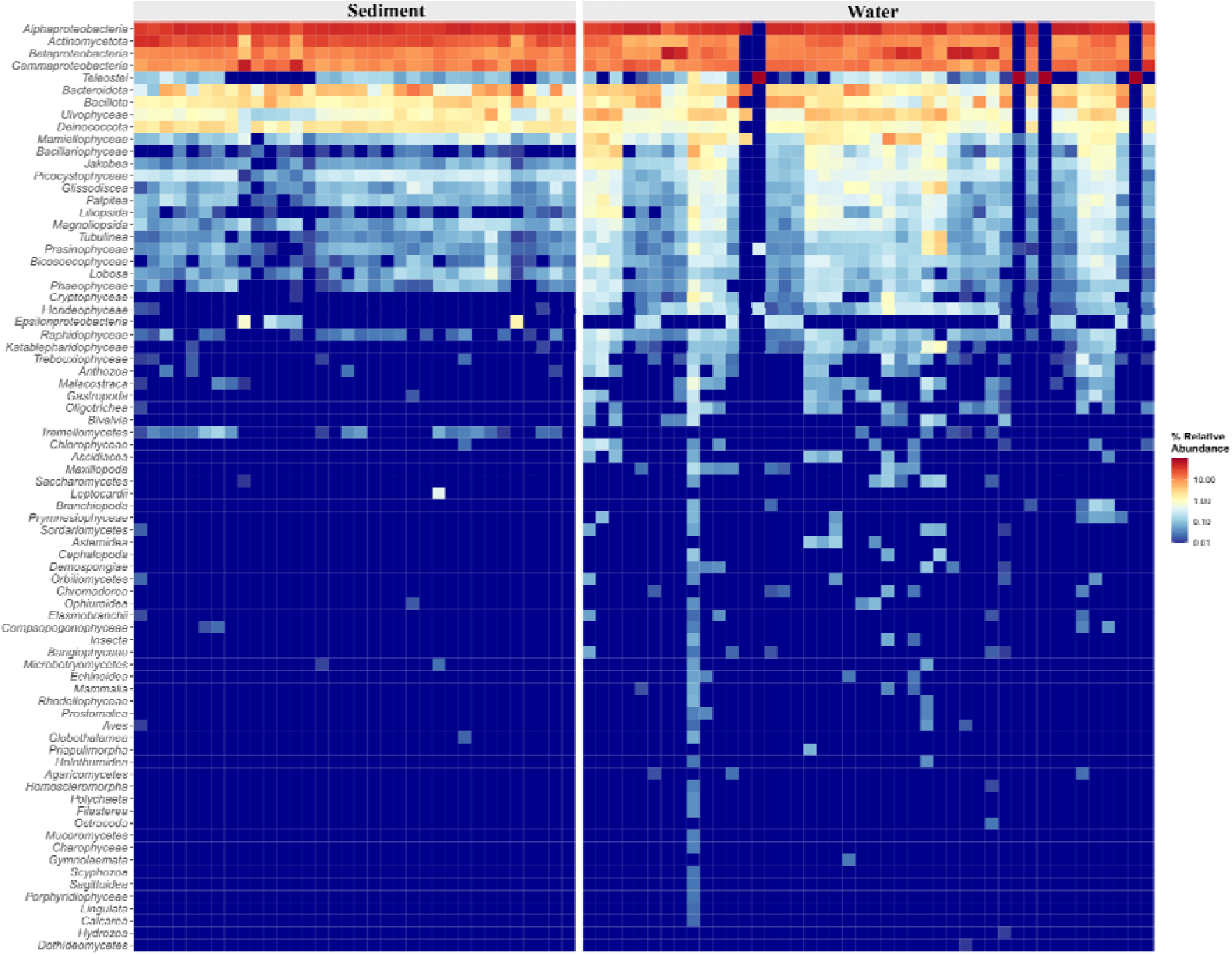
Heatmap showing the relative abundance of assembled contigs across taxonomic classes identified in the eDNA dataset. Relative abundance values range from 0.01 to 20, with higher values represented in brick red and lower values in deep blue.

The number of families varied accross kingdoms. The kingdom Animalia was the most diverse, spanning 189 families, dominated by *Gobiidae*, alongside other reef- and pelagic-associated fish and invertebrate families (Table S6). Bacteria comprised 29 families, with *Eubacteriales* being the most dominant, whlie Plantae comprised 37 families, with *Gracilariaceae, Ulvaceae,* and *Solieriaceae* being the most represented. Rhizophoraceae, a known mangrove family, was also detected. *Chromista* was represented by 31 families, and largely dominated by *Bacillariaceae* (Table S6). Fungi were less diverse, comprising 12 families, with *Debaryomycetaceae* being the most represented. Protozoa was the least diverse, with only six families represented (Table S6).

### Spatial patterns of eDNA-derived OTU assemblages across samples and sites

OTU composition, richness and diversity were comparable across all sampled sites and sample types (Fig. 7a). Comparison of OTUs between sites showed no significant differences in richness (Chao1: χ² = 9.54, df = 16, p = 0.890), community diversity and distribution (Shannon: χ² = 14.00, df = 16, p = 0.599), nor in dominance (Simpson: χ² = 21.66, df = 16, p = 0.154). Comparison between sample types showed significant variation in OTUs diversity (Figure 7a). OTU richness was higher in water samples than in sediment (Chao1: χ² = 25.77, df = 1, p < 0.001; Dunn’s tests: Z = −5.08, p < 0.001, Fig. 7b). Water samples also had higher OTU diversity and distribution compared to sediment (Shannon: χ² = 6.09, df = 1, p = 0.014; Dunn’s test: Z = −2.47, p = 0.014, Fig. 7c), while OTU dominance was insignificant accross sample types (Simpson: χ² = 1.25, df = 1, p = 0.263; Dunn’s test: Z = 1.12, p = 0.263).

**Figure 7.**
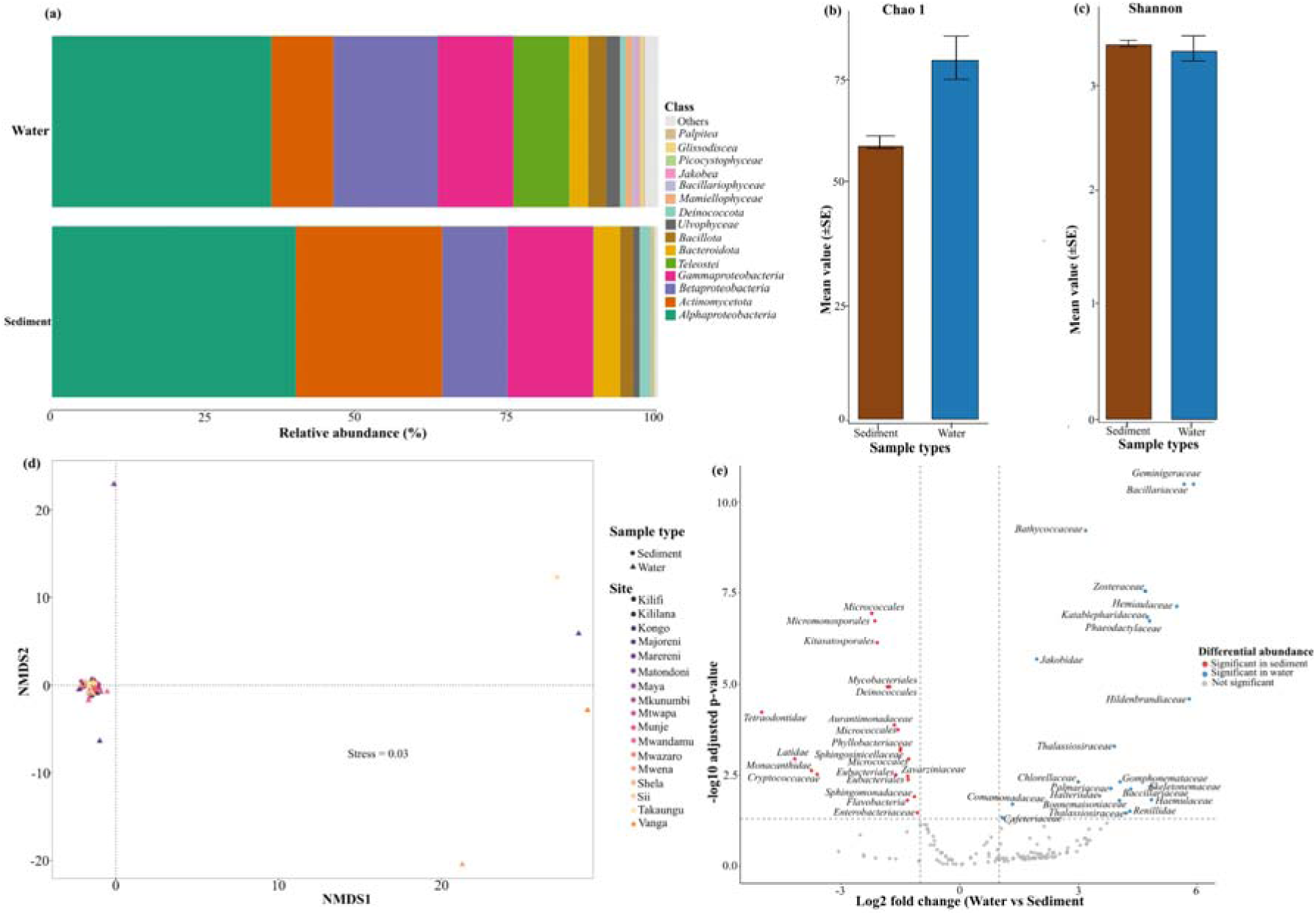
Operational taxonomic unit (OTU) diversity and community composition. (a) Comparison of alpha diversity among study sites. (b) Alpha diversity (Chao1) across sample types. (c) Alpha diversity (Shannon) across sample types. (d) Non-metric multidimensional scaling (NMDS) ordination showing compositional differences in OTUs detected between sample types. (e) Differentially abundant taxa between water and sediment samples.

Evaluation of variation in community assemblage showed overlap in OTU composition within sites (PERMANOVA: R² = 0.33, F = 14.66, p = 0.001) and between sample types (PERMANOVA: R² = 0.16, F = 1.85, p = 0.001). NMDS ordination based on Jaccard distances showed clear structuring of OTU assemblages across sites and sample types (Fig. 7d), with water samples having greater dispersion (stress = 0.03). Differential abundance analysis resolved OTU-level differences between water and sediment samples (Fig. 7e). Sediment samples showed significantly higher bacterial abundance than water samples, with most significant OTUs belonging to the kingdom bacteria (Fig. 7e). Contigs assigned to fungal and some fish OTUs were also significantly abundant in sediment samples (Fig. 7e). Water samples had significant abundance of OTUs belonging to phytoplankton, algae, and free-living aquatic microbes, diatom, microalgal, green and red algal in water samples than in sediment samples (Fig. 7e).

### Complementarity of literature records and snapshot eDNA detections

Literature review and eDNA survey together recorded a cumulative total of 488 families. The literature survey documented 251 families, of which 183 were unique, while the eDNA survey detected 305 families, with 238 unique to eDNA. The overlap between the two datasets included 67 families, comprising 49 fish families (Teleostei), three bird families (Ardeidae, Charadriidae, Scolopacidae), four crustacean families (Grapsidae, Ocypodidae, Portunidae, Penaeidae), six bivalve families (Chthamalidae, Muricidae, Naticidae, Ostreidae, Strombidae, Potamididae), and one red algae family (Gracilariaceae) (Figure 8). Among *teleostei*, eDNA detected 33 families absent from the literature survey, while the literature survey recorded 25 families not captured by eDNA. eDNA also detected additional non-vertebrate taxa, macroalgae, echinoderms, mollusks, crustaceans, fungi, and protists, which were not adequately represented in the literature. *Aves* and *Nematoda* were better documented in the literature survey, although eDNA detected Stercorariidae family, which was absent from the literature records.

**Figure 8:**
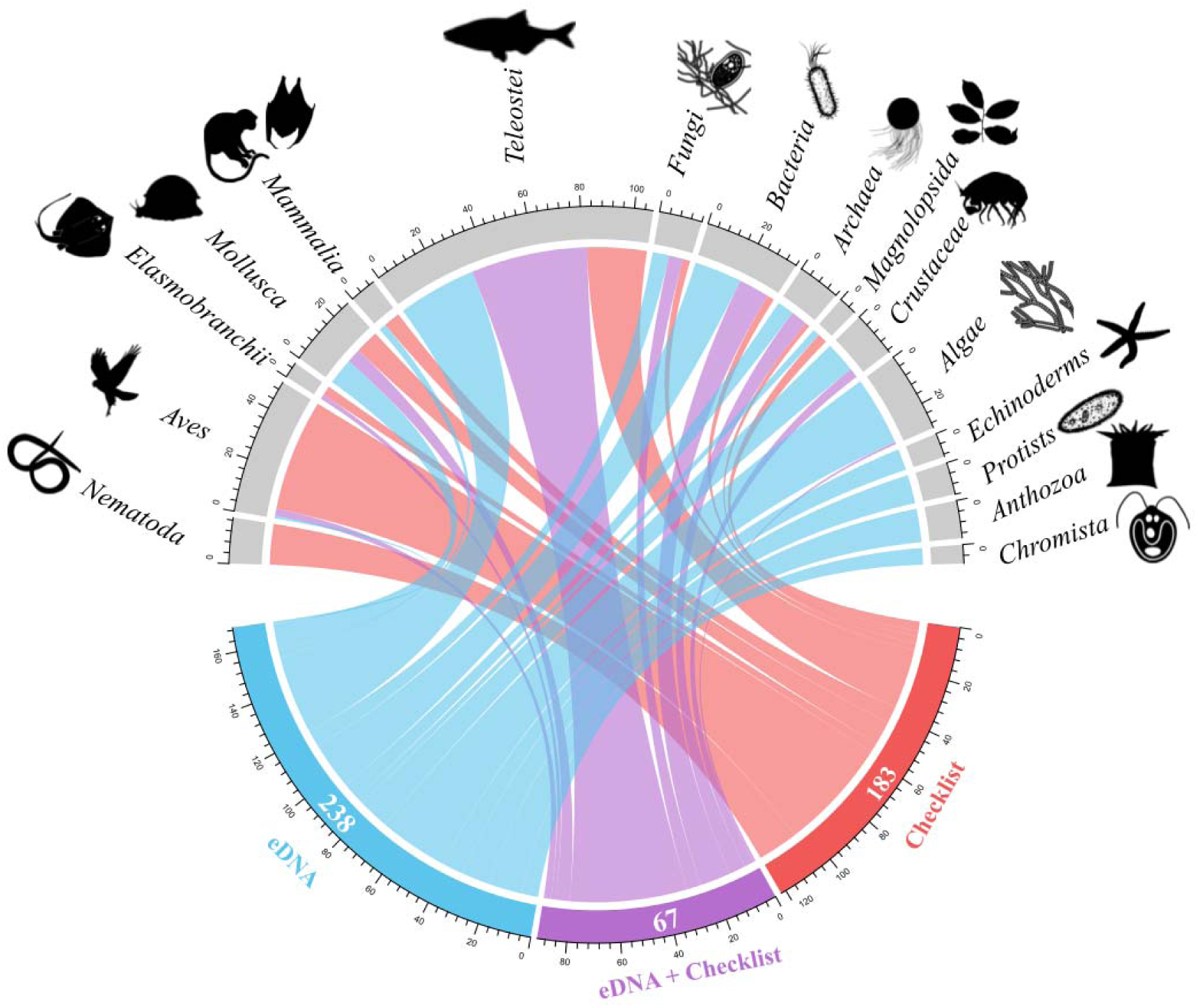
Chord diagram illustrating taxonomic overlap between eDNA detections and literature review. The diagram represents 488 taxonomic families, of which 305 were identified by eDNA and 251 from the literature review. Chords denote taxa unique to eDNA (238), common to both (67), and unique to the literature review (183). The upper panel displays taxa at different classification levels.

## Discussion

Comprehensive biodiversity assessments are essential for informing management strategies (Kopnina *et al*., 2024; Tobias *et al*., 2025). Despite increasing attention to mangrove rehabilitation and conservation, significant gaps in biodiversity surveys persist, limiting the understanding of biodiversity dynamics. Evaluating how mangrove biodiversity has been studied historically, and whether approaches such as eDNA can help address these gaps, is therefore key to developing a complete understanding of biodiversity in mangroves and supporting evidence-based management. As a proof of concept, this study demonstrates that mangrove biodiversity studies in Kenya are inadequately structured, with no clearly defined or standardized criteria guiding assessments. Research efforts have been spatially uneven (Fig. 4, Fig. 3d), temporally inconsistent (Fig. 2a), and disproportionately focused on a limited subset of taxa (Fig. 3c, Fig. 4) and sites (Fig. 3d, Fig. 4). This pattern mirrors broader trends reported across many ecosystems in developing countries within the tropics, where biodiversity studies have remained fragmented and biased (Titley *et al*., 2017; Tydecks *et al*., 2018; Guerrero-Casado & Monge-Nájera, 2021; Moreno *et al*., 2023). While eDNA metagenomics generates high-throughput data capable of characterizing mangrove biodiversity and offering an alternative approach to addressing existing data gaps (Wee *et al*., 2023; Panda *et al*., 2025), the findings of this study reveal a more nuanced reality, highlighting current limitations in the resources and infrastructure required to reliably streamline application of eDNA for mangrove biodiversity studies in Kenya.

### Review of historical biodiversity data

The literature review revealed variation in biodiversity assessment efforts across mangrove forests in Kenya (Fig. 4). Since 1990, research has been concentrated in Gazi Bay, Mida Creek and Tudor Creek, with comparatively reduced effort in other forest blocks (Fig. 3d, Fig. 4). Although these forest blocks represent less than 6% of total mangrove forest cover in Kenya (GoK, 2017), they account for 63.15% (Fig. 4) of all the documented biodiversity studies, suggesting their importance in mangrove ecological research. In contrast, Lamu, despite containing nearly 60% of mangrove forest cover in Kenya (GoK, 2017; Kairo *et al*., 2021), had no biodiversity survey records in any of the reviewed studies. This imbalance indicates the poor state of biodiversity studies and is likely attributable to established research priorities focused on ecological restoration and blue carbon (Kairo *et al*., 2018, 2021, 2025), inadequate resources to conduct regular comprehensive biodiversity assessments (Siddig, 2019), and insecurity (Laher, 2011). The higher number of studies in Gazi Bay is not coincidental (Fig. 3d). For several years, the site has served as a hub for mangrove research, particularly in restoration ecology, supported by initiatives such as the Mikoko Pamoja project (Gallin *et al*., 1989; Kinya *et al*., 2024; Locatelli *et al*., 2014). The presence of such projects likely attracted sustained funding and scientific attention, facilitating parallel biodiversity assessments aimed at establishing baselines or evaluating restoration outcomes (Crona *et al*., 2006; Kirui *et al*., 2008; Mutua *et al*., 2011).

Similarly, Mida Creek, where a comparatively higher number of studies were identified (Fig. 3d, Fig. 4), is an important habitat for resident and migratory bird species, serving as a key stopover along the Western Indian Ocean flyway (Pearson *et al*., 2002). Its prominence as a birding, conservation, and ecotourism destination has facilitated relatively consistent ornithological surveys (Moragwa *et al*., 1996; Nussbaumer *et al*., 2024; Seys *et al*., 1995). Tudor Creek, on the other hand, has been a model system for examining the impacts of urbanization on mangrove structure and functioning (Bosire *et al*., 2014; Mohamed *et al*., 2016), which likely explains the number of biodiversity-related studies identified (Fig. 3d). For most mangrove sites, biodiversity assessments were conducted only once, with no evidence of sustained or systematic monitoring (Fig. 3d, Fig. 4). Given the structural complexity of mangrove forests and the logistical challenges associated with conventional biodiversity surveys, this likely reflects limited financial resources and a shortage of trained personnel capable of conducting regular assessments (Stephenson et al., 2020). Nearly 90% of mangrove forests in Kenya lacked published biodiversity data, representing a substantial knowledge gap.

The number of biodiversity studies has remained modest, and the frequency of surveys partially aligns with general recommendations for periodic biodiversity monitoring (England *et al*., 2021; Gatt *et al*., 2022). However, this level of effort is insufficient for establishing ecological baselines or for evaluating the long-term outcomes of restoration and management interventions (Dalton *et al*., 2023; Sunkur *et al*., 2024). The reviewed studies lacked a systematic framework that would enable comparative analyses across sites and time (Fig. 3d). Effective ecological monitoring typically requires consistent, long-term surveys conducted across broad environmental gradients, targeting multiple taxa and sustained over time to detect trends and understand underlying ecological processes (Brown *et al*., 2023; Harvey *et al*., 2020). The reviewed studies showed no evidence of wide spatial coverage, follow-up assessments of consistent species, integration of results into national biodiversity databases, or linkage to ongoing national or global monitoring initiatives. This suggests that most of the studies were short-term and project-specific, often conducted for academic purposes, and designed to meet immediate objectives rather than to contribute to a structured, long-term understanding of mangrove biodiversity dynamics.

The disproportion in the taxa identified, where the classes *Teleostei, Aves, Chromadorea*, and *Malacostraca* accounted for 85.4% of all recorded taxa (Fig. 4), suggests the importance of these groups in socioeconomic and ecological roles. The dominance of *Teleostei* (ray-finned fish), comprising more than half of the recorded taxa (56.1%; Fig. 4), points to a long-standing research emphasis on this group. Teleostei are a source of livelihood for coastal communities in Kenya, generating income for artisanal fisherfolk and supporting commercial fisheries (Ahmed *et al*., 2022; Morara *et al*., 2015; Samoilys *et al*., 2017). Under the Blue Economy agenda, the Government of Kenya has identified fish resources as strategic assets for economic growth. Significant investments have been made in data collection and research to support sustainable fisheries management and to explore opportunities for increasing fish production and productivity (Njiru *et al*., 2021). Additionally, *Teleostei* have been widely used as indicator species for assessing the role of mangroves as breeding and nursery habitats (Gajdzik *et al*., 2014; Hernández-Mendoza *et al*., 2022; Igulu *et al*., 2014; Lefcheck *et al*., 2019; Pinna *et al*., 2023). Their high representation in the survey records, therefore, aligns with research priorities focused on evaluating the ecological importance of mangroves and their relevance to livelihood ventures and fisheries management. Similarly, *Aves, Chromadorea*, and *Malacostraca* were also represented in relatively high proportions (Fig. 4), consistent with their established use as indicator taxa for evaluating ecosystem health and the success of restoration and rehabilitation initiatives (Mutua *et al*., 2011; Nudi *et al*., 2007; Smits & Fernie, 2013). *Aves* were mainly documented in studies conducted in Mida Creek and the Sabaki estuary, with few records in other forest blocks (Fig. 3a). This pattern is most likely attributed to the well-developed bird observer networks, citizen science initiatives, and long-standing monitoring traditions, which have facilitated consistent record collection in many regions globally (Seys *et al*., 1995). Taxa such as Chromadorea and Malacostraca were predominantly documented in Gazi Bay (Fig. S3), where they were mainly used in ecological monitoring following restoration and management interventions. The narrow taxonomic focus of previous mangrove biodiversity studies has yielded only marginal increases in newly detected taxa over time (Fig. 3b), suggesting substantial gaps in taxonomic coverage remain. This pattern likely reflects established study priorities and methodological limitations rather than a true lack of biological diversity, indicating that mangrove forests in Kenya remain considerably undercharacterised.

### Environmental DNA findings

The large number of reads sequenced in this study demonstrates the potential of eDNA to detect broad-scale biodiversity signals in mangrove systems. From over 5 million assembled contigs, only 1.65% were assigned to OTUs at high sequence similarity. This notable small number of assigned contigs can be attributed to stringent quality thresholds used and gaps in the reference databases; a pattern consistently reported in metagenomic and metabarcoding eDNA studies (Macé *et al*., 2022; Mathon *et al*., 2021; Mwamburi *et al*., 2023; Patin *et al*., 2025; Serivichyaswat *et al*., 2025). Following low abundance and occurrence filtering, the retention of 2,330 OTUs represents a conservative and ecologically meaningful subset of the data. Such filtering approaches are widely applied in eDNA studies to minimize the influence of sequencing artefacts, index hopping, and sporadic low-frequency detections, while retaining OTUs that are more likely to reflect true biological signals and recurrent taxa in the environment (Evans *et al*., 2017; Peixoto *et al*., 2021).

The detection of OTUs from multiple ecosystems (Fig. 5a) suggests that the mangrove system is highly connected to adjacent habitats through hydrological exchange processes, where it functions as a filter and a conduit for DNA derived from surrounding environments. Previous studies have demonstrated that tidal flushing, riverine discharge, surface runoff, and aerial transport can introduce allochthonous DNA into coastal habitats (Andruszkiewicz *et al*., 2019; Deiner *et al*., 2016; Shogren *et al*., 2017). Although this complexity complicates the interpretation of resident mangrove assemblages, it simultaneously provides insight into carbon sources within mangrove ecosystems and the degree of ecological connectivity with adjacent habitats, information that may be valuable for integrated management and conservation planning. Cross-referencing with the regional species checklists showed that nearly two-thirds of detected OTUs were classified as “unexpected”. This pattern should not be interpreted solely as evidence of false positives or biological implausibility. Numerous studies have shown that regional biodiversity inventories remain incomplete and taxonomically biased toward charismatic and economically important taxa (Mora *et al*., 2011). Similarly, genetic reference databases are unevenly populated, with microorganisms, invertebrates, algae, and cryptic or understudied taxa being underrepresented (McGee *et al*., 2019; Weigand *et al*., 2019). As a result, many eDNA-derived OTUs cannot be confidently matched to existing regional records, even when they represent genuine components of local or adjacent biodiversity. “Unexpected” detections may therefore correspond to undocumented regional diversity, transient taxa transported from connected ecosystems, or lineages that remain taxonomically unresolved due to gaps in reference sequences. The persistence and transport of eDNA through hydrological processes can also introduce signals from non-resident organisms or past communities, further complicating direct comparisons with static regional species checklists (Deiner *et al*., 2017).

After restricting analyses to marine, brackish, and ubiquitous taxa, the retained OTUs still had taxonomic coverage, spanning 76 classes and 305 families. The dominance of bacterial contigs relative abundance is consistent with the ubiquity, high biomass, and rapid turnover of microbial communities in mangrove environments. Classes such as Alphaproteobacteria, Gammaproteobacteria, and Actinomycetes are well known to have an integral role in nutrient cycling, organic matter decomposition, and biogeochemical processes in coastal wetlands, supporting the ecological relevance of their high representation (Liang *et al*., 2023). In contrast, other groups, such as plants, fungi, and chromists, showed lower relative contig abundance, likely indicating biological differences in DNA shedding and methodological biases related to DNA extraction efficiency and genome copy number. The dominance of the class *Teleostei* among OTUs mirrors long-standing research and database biases rather than true community composition. *Teleosts* are the most studied vertebrates due to their economic importance and use as indicator species (Ahmed *et al*., 2022; Njiru *et al*., 2021; Samoilys *et al*., 2017). As a result, they are disproportionately represented in genetic reference databases, facilitating higher confidence taxonomic assignments. Other animal groups, although present, are comparatively underrepresented, providing evidence on how reference availability shapes apparent biodiversity patterns in eDNA studies. Similar biases are evident in the bacterial and plant datasets, where well-characterized lineages dominate assignments.

Patterns of OTU richness and diversity across sites revealed broadly comparable biodiversity levels, suggesting that mangrove-associated communities may be relatively homogeneous at the scale examined, or that high connectivity among sites promotes similar assemblages. The lack of strong site-specific dominance supports the idea of shared regional species pools. In contrast, significant differences between water- and sediment-derived eDNA illustrate the importance of sampling substrate in biodiversity inference (Fig. 7b, Fig. 7c). Water samples captured higher richness and diversity, likely indicating more recent, mobile, and spatially integrated DNA signals, whereas sediments may preserve more localized and temporally integrated assemblages (Fig. 7e). Multivariate analyses nonetheless revealed significant heterogeneity in community composition across sites and sample types, despite overlapping assemblages. The detection of specific indicator families, ecologically and economically important fish, crustaceans, and microbial lineages, demonstrates that eDNA is sensitive enough to detect subtle shifts in assemblage structure. These patterns likely arise from variation in habitat characteristics, hydrodynamic regimes, and localized human activities, indicating the added value of eDNA in resolving fine-scale ecological variation that is often difficult to detect using conventional techniques.

### Integrating literature and environmental DNA results for comprehensive view of mangrove biodiversity

A look into the literature review and eDNA-derived detections at the family level revealed considerable differences in taxonomic composition, consistent with fundamental methodological contrasts between conventional survey approaches and eDNA-based surveys (Fediajevaite *et al*., 2021; Iacaruso *et al*., 2025). eDNA detected a higher number of unique OTUs per family than those reported in historical checklists (Fig. 8), in line with its high sensitivity and capacity to detect a broad spectrum of taxa across multiple kingdoms (Deiner *et al*., 2017; Thomsen & Willerslev, 2015). Rather than indicating disagreement between datasets, these patterns indicate that historical checklists and eDNA surveys capture complementary dimensions of biodiversity rather than overlapping inventories. Literature records largely represent accumulated data defined by targeted sampling designs, taxon-specific expertise, and long-standing research and management priorities, which tend to favor conspicuous, economically important, or morphologically distinct taxa (Bohmann *et al*., 2014). In contrast, eDNA detected a snapshot of biodiversity from DNA traces from a wide range of organisms irrespective of their detectability in the field, thereby extending taxonomic coverage beyond the reach of conventional approaches. The minimal overlap observed between the two datasets is best explained by differences in detection pathways rather than inconsistencies in underlying biodiversity patterns. Taxa shared between both sources were dominated by well-studied groups with extensive sampling histories and well-curated reference databases, whereas families detected exclusively by eDNA, particularly among algae, invertebrates, fungi, and protists, demonstrate the capacity of the technique to recover understudied and invertebrate components of mangrove ecosystems that are poorly represented or absent in historical records (Pawlowski *et al*., 2018; Cordier *et al*., 2021). Reduced detection efficiency of eDNA for certain groups, such as birds and nematodes, aligns with known biological and methodological constraints such as low DNA shedding rates, and rapid degradation in aquatic environments (Ushio *et al*., 2017; Deiner *et al*., 2019). The detection of additional fish families by eDNA may indicate historical under-sampling, improved sensitivity of the eDNA technique, or recent changes in community composition.

These findings should nevertheless be interpreted in light of several study limitations that inform priorities for future research. The synthesis of reviewed biodiversity information relied exclusively on accessible published documents, which likely provides an incomplete representation of mangrove biodiversity in Kenya, as unpublished institutional datasets, online databases, and archived hard-copy records were not systematically examined. In parallel, a substantial proportion of eDNA-derived OTUs could not be confidently assigned to known taxa or matched to regional species inventories, revealing gaps and taxonomic biases in available reference databases. Interpretation of eDNA signals was also complicated by the dynamic nature of mangrove ecosystems, which accumulate genetic material from adjacent habitats through tidal fluxes, runoff, and sediment transport, making it difficult to distinguish resident organisms from exogenous DNA (Teixeira *et al*., 2023; Zainal Abidin *et al*., 2022). Methodological factors, such as variation in DNA extraction efficiency, differences in DNA shedding among taxa, substrate-dependent DNA persistence, and the choice of database, may also have influenced detection probabilities and shaped observed community patterns. Although marker-based sequencing approaches such as COI, 18S rRNA, ITS, 16S rRNA, and 12S rRNA are widely used for taxa resolution and generally appropriate, they are susceptible to amplification biases (Teixeira *et al*., 2023; Zainal *et al*., 2022), are prone to misidentifying organisms when taxa are resolved to lower ranks (Kislichkina *et al*., 2025), and are costly when multiple taxa are targeted. Addressing these challenges will require coordinated efforts to expand biodiversity syntheses beyond the accessible published literature, develop region-specific genetic reference libraries supported by voucher-based barcoding, and integrate eDNA in targeted conventional surveys using standardized, replicated sampling designs across multiple substrates and seasons. Long-term archiving of raw sequence data and physical samples will facilitate retrospective analyses as analytical methods and reference resources advance, reinforcing the potential of eDNA as a suitable tool for mangrove biodiversity research and monitoring in Kenya.

## Conclusion

Biodiversity studies in mangrove forests in Kenya have progressed slowly, and the available information on taxonomic and spatiotemporal coverage remains limited. Most documented surveys have been concentrated in Gazi Bay, Mida Creek and Tudor Creek, which may reflect historical sampling biases, leaving extensive areas of mangrove forest insufficiently assessed. Recorded taxa are also unevenly represented, with *Teleostei, Aves, Chromadorea* and *Malacostraca* comprising a large proportion of the species reported. This pattern suggests that the full coverage of biodiversity present in mangroves is yet to be characterized. The use of eDNA revealed a wider spectrum of taxa, ranging from prokaryotes to eukaryotes, and detected a level of diversity that exceeds what has been documented through conventional surveys. This demonstrates the utility of eDNA in expanding biodiversity assessment in mangroves. However, mangroves are heterogeneous environments with genetic material from multiple sources, making it challenging to determine which detections reflect resident species, especially when using metagenomic studies. The absence of regional genetic reference databases also limits accurate taxonomic assignment, resolution and variation in DNA persistence among sample types may influence interpretation. Effective implementation of eDNA for biodiversity assessment in mangroves in Kenya will require resolving these basic challenges to improve confidence in the results and strengthen its application as a survey tool. Addressing these issues will also support the gradual incorporation of eDNA into existing monitoring frameworks. At present, eDNA is most effective when used together with conventional field approaches to help verify and validate detections.

## Acknowledgement

This study was funded by Kenya Marine and Fisheries Research Institute, Plan International and Wetlands International Eastern Africa. We are grateful to Laureen Amondi, Mathews Wafula, Jane Mutheu, Sarah Anyango, Isaac Mondestor, Diana Chepngetich, Oliver Ogola, and Miriam Vihenda for their invaluable support in fieldwork, sample processing, and literature review. Special thanks to Jackline Olando and Christine Mwigwi of KMFRI for their assistance with laboratory logistics, and Belinder Ochillo for preparing the map.

## Data availability

All environmental DNA data generated in this study will be deposited in the National Center for Biotechnology Information (NCBI) repository prior to publication.

## Authors contribution

**OFO** conceptualize the idea, data curation, data analysis, Writing the original draft and editing, investigation, methodology, and visualization. **JO** - Conceptualize the idea, resource mobilization, writing the original draft, and editing. **SM** - Conceptualize the idea, data analysis (Bioinformatics), writing-original draft and editing **ZM** - Data curation, data analysis, writing the original draft, and editing, **EO** - Data curation, writing the original draft, and editing, **SC** - Data curation, writing and editing the original draft, **CL** Data curation, writing and editing the original draft.

