## Supplementary material for "State of mangrove biodiversity assessment in Kenya and the prospect of environmental DNA in strengthening surveys": Fig. S1

**Corresponding Authors***

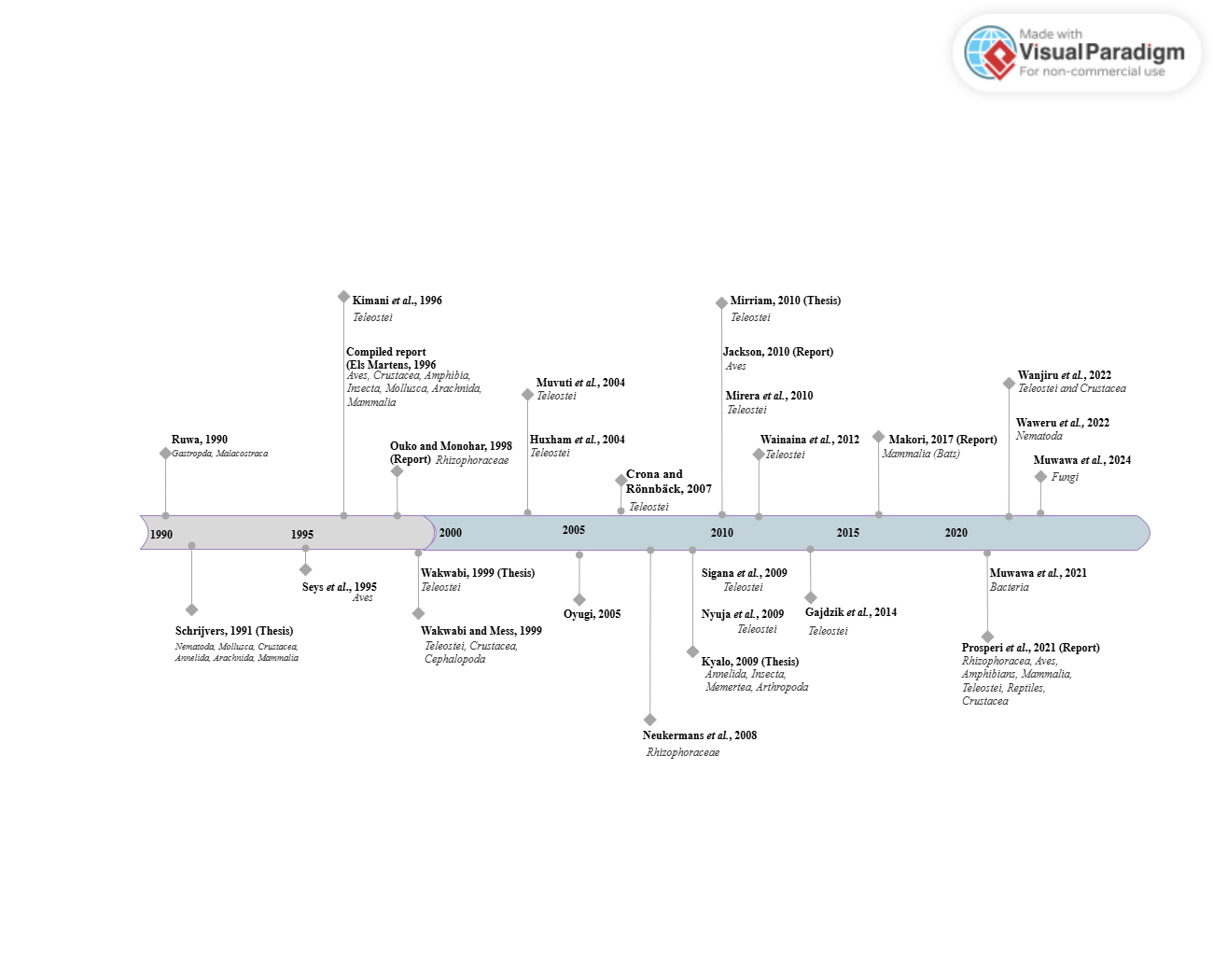

**Figure S1:** Schematics illustrating studies published on mangrove biodiversity assessments in Kenya from 1990 to 2024

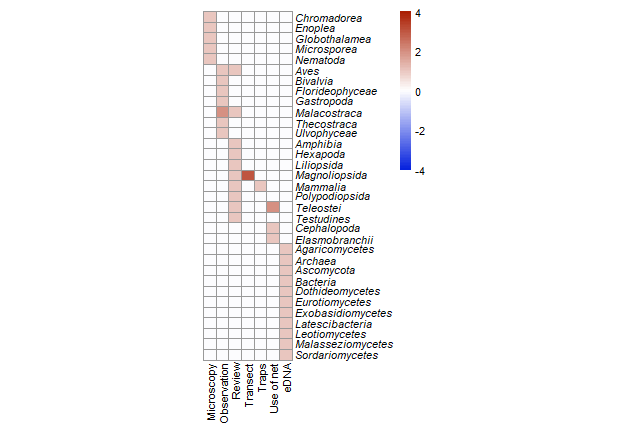

**Figure S2**: Survey techniques used to assess mangrove biodiversity over the years. eDNA approaches were predominantly used to target microbial communities, while conventional techniques were primarily used to survey macro-organisms.

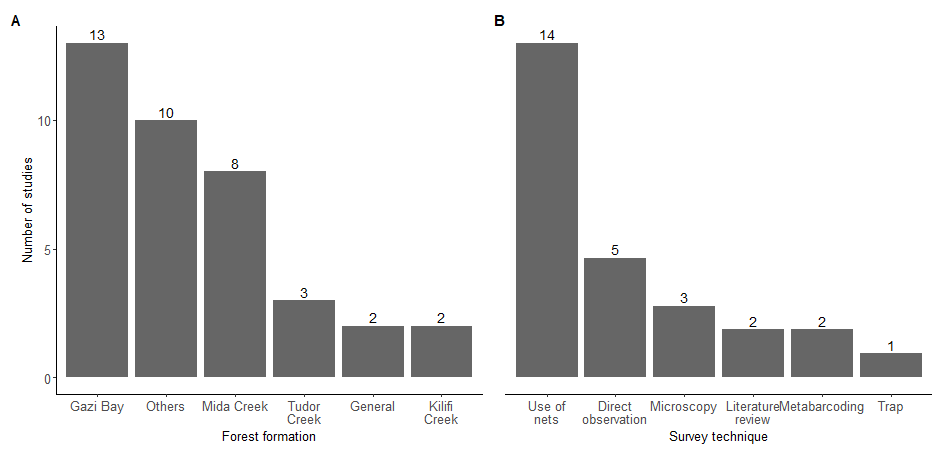

**Figure S3**: A) Number of studies conducted in each mangrove forest. "Other" includes Bamburi, Kanamai, Mkomani, Malindi, Sabaki, Shimoni, Mtwapa Creek, Ungwana Bay, and Vanga, each of which was surveyed only once. B) Number of survey techniques used across studies.

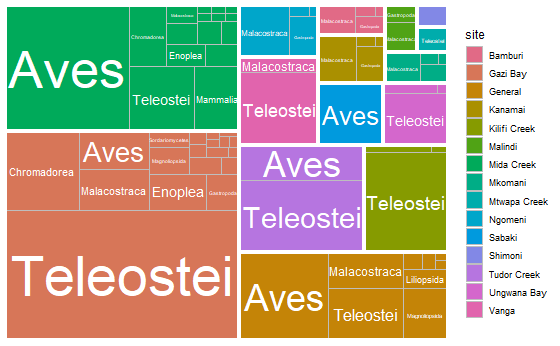

**Figure S4**: Treemap showing the proportional representation of taxa surveyed in individual mangrove forests and across general studies over the years.

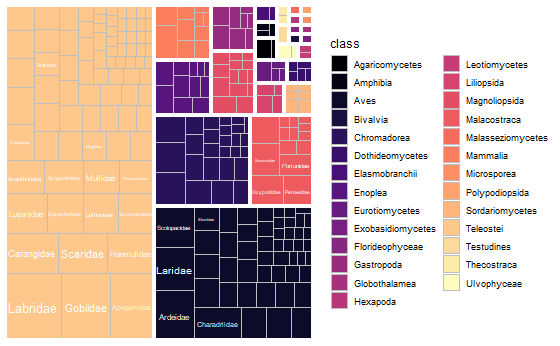

**Figure S5**: Treemap showing the relative proportion of families within each class identified in the review.

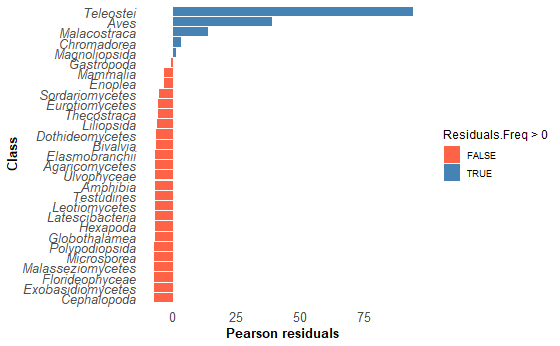

**Figure S6:** Pearson correlation illustrating the taxonomic survey bias of biodiversity surveye in mangrove forests in Kenya. Taxa belonging to the classes *Teleostei, Aves, Malacostraca, Magnoliopsida,* and *Chromadorea* were disproportionately studied relative to other groups during the period 1990–2024.

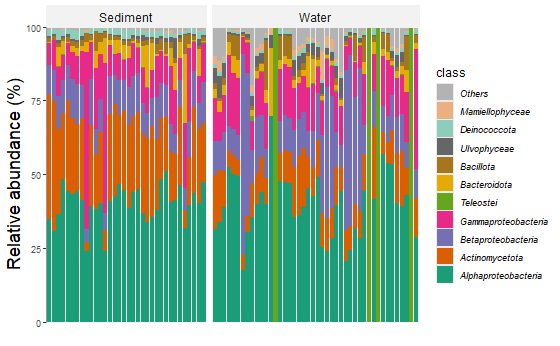

**Figure S7**: Relative abundance of reads across phyla identified using an environmental DNA approach.
